# Tracking infection dynamics at single-cell level reveals highly resolved expression programs of a large virus infecting algal blooms

**DOI:** 10.1101/757542

**Authors:** Chuan Ku, Uri Sheyn, Arnau Sebé-Pedrós, Shifra Ben-Dor, Daniella Schatz, Amos Tanay, Shilo Rosenwasser, Assaf Vardi

## Abstract

Nucleocytoplasmic large DNA viruses have the largest genomes among all viruses and infect diverse eukaryotes across various ecosystems, but their expression regulation and infection strategies are not well understood. We profiled single-cell transcriptomes of the worldwide-distributed microalga *Emiliania huxleyi* and its specific coccolithovirus responsible for massive bloom demise. Heterogeneity in viral transcript levels detected among single cells was used to reconstruct the viral transcriptional trajectory and to map cells along a continuum of infection states. This enabled identification of novel viral genetic programs, which are composed of five kinetic classes with distinct promoter elements. The infection substantially changed the host transcriptome, causing rapid shutdown of protein-encoding nuclear transcripts at the onset of infection, while the plastid and mitochondrial transcriptomes persisted to mid- and late stages, respectively. Single-cell transcriptomics thereby opens the way for tracking host-pathogen infection dynamics at high resolution within microbial communities in the marine environment.

## Main text

Nucleocytoplasmic large DNA viruses (NCLDVs) are the largest viruses known today in both genome and virion size. They have been found in most major lineages of eukaryotes across diverse habitats (*1*–*4*), especially in the marine environment (*5*, *6*). Among the NCLDVs of special ecological importance are members of the family *Phycodnaviridae* that infect a wide range of key algal hosts (*1*). These include the cosmopolitan calcifying eukaryotic alga *Emiliania huxleyi* (Haptophyta), which forms massive annual blooms in the oceans that have a profound impact on the carbon and sulfur biogeochemical cycles (*7*). *E. huxleyi* blooms are frequently terminated by a large dsDNA virus — EhV (*8*), which enhances nutrient cycling and carbon export to the deep ocean (*9*, *10*). This host-virus model provides a trackable system for understanding viral life cycle strategies and host responses.

High-throughput bulk RNA sequencing (RNA-seq) has been used for whole-genome expression profiling of NCLDVs during infection, shedding light on gene prediction, transcript structure, and changes in metabolic pathways (*11*–*13*). However, bulk RNA-seq profiles average gene expression levels across many cells, whereas infection states can be variable among single cells. To overcome this limitation, single-cell RNA-seq (scRNA-seq) approaches have been developed to probe the transcriptomes of individual cells in a highly parallel manner. These methods have revolutionized our understanding of various developmental and immunological processes (*14*, *15*), including host-virus interactions in mammalian systems (*16*, *17*).

Here we employed scRNA-seq to study EhV infection of *E. huxleyi* at the single-cell level, in order to characterize the temporal dynamics and regulation of viral and host transcriptomes. *E. huxleyi* CCMP2090 cultures were infected with the lytic virus EhV201 (*13*, *18*) (Fig. 1A, Table S1, and fig. S1). Individual cells were isolated into 384-well plates by fluorescence-activated cell sorting (FACS) and processed using the MARS-seq protocol (*19*), which uses cellular barcodes to tag RNA molecules from individual cells and enables transcript quantification by using unique molecular identifiers (UMIs). By mapping reads to the reference *E. huxleyi* transcriptome and to the EhV201 genome, we were able to profile both viral and host transcripts in each individual cell.

**Fig. 1.**
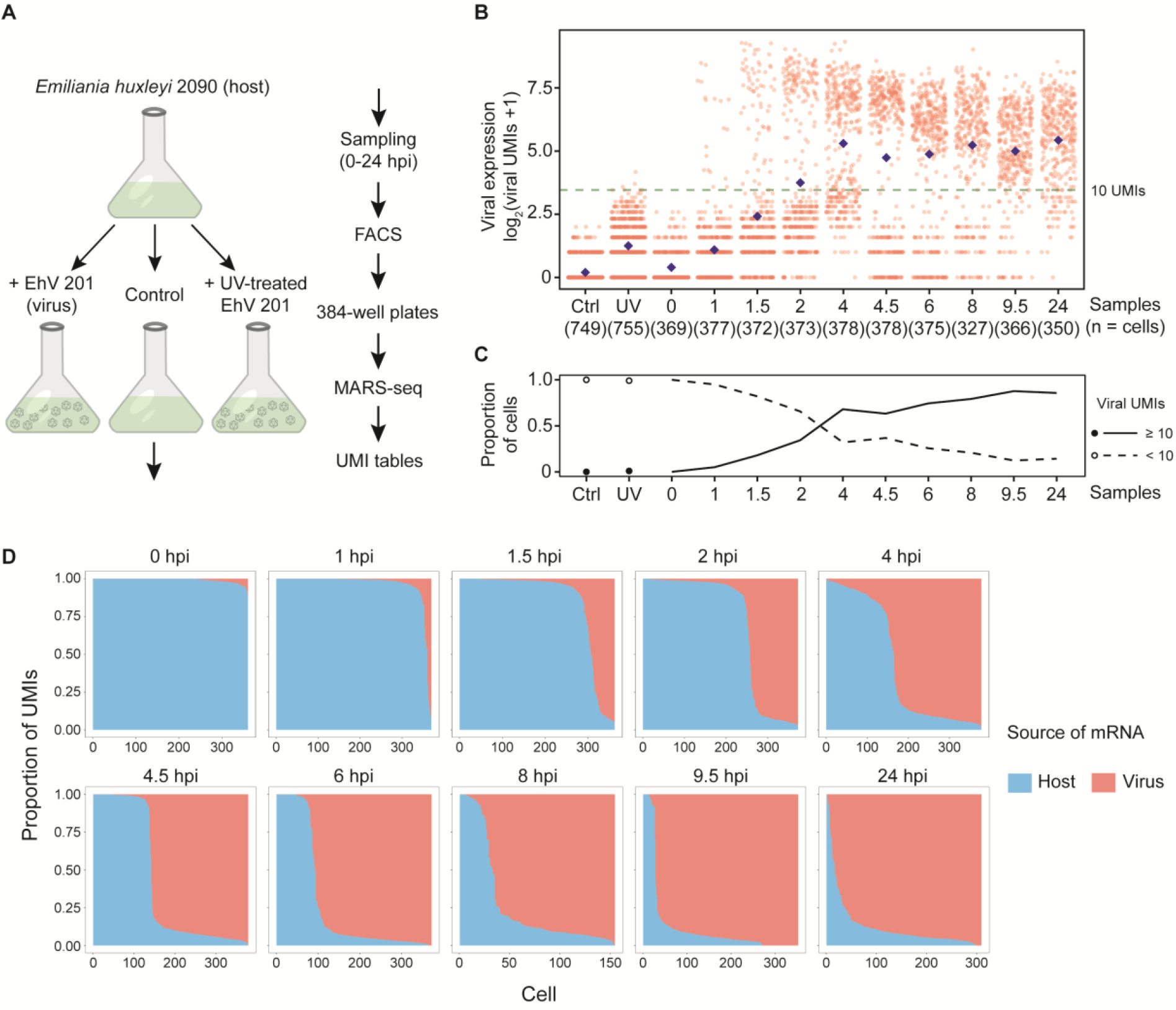
Transcriptome profiling of EhV infection in *E. huxleyi* at single-cell resolution. (**A**) The experimental setup for using single-cell RNA-seq to study EhV infection of *E. huxleyi*. Individual cells were isolated by FACS during a time course of infection. The single-cell transcriptomes were sequenced using the MARS-seq protocol, which allows absolute quantification of viral and host transcripts by tagging each individual cell and each transcript by cellular barcodes and unique molecular identifiers (UMIs), respectively. (**B**) Highly heterogeneous total viral UMI counts among single cells across samples. Each red dot represents an individual cell and the blue diamonds the average expression of all cells at each time point. A threshold of 10 UMIs (fig. S3) highlights the bimodality of overall viral expression and excludes cells only with low- or noise-level viral UMIs, as in mock infection by UV-deactivated EhV (UV; 4 and 6 hpi) and control (Ctrl) samples. (**C**) The proportion of cells with at least 10 viral UMIs (active viral expression) increased rapidly during early hours of infection and approached 90% at 9.5 hpi. (**D**) Cliff diagrams represent the relative mRNA levels of host and virus across the infection time course. The cells (columns) are sorted by the relative proportions of host and viral mRNA, showing the sharp decrease in host mRNA relative to viral mRNA (“cliff”) and the within-population heterogeneity through time. hpi: hours post infection.

We compared the total expression levels of all viral genes in 5,179 cells from infected cultures, control cultures, and cultures mock-infected by UV-inactivated viruses (Fig. 1B). While the average viral gene expression increased over time of infection, high variability in viral expression levels between single cells was observed despite a high virion-to-cell ratio (multiplicity of infection 5:1) that was used to reduce the possible variation due to encounter rates. Throughout the first 10 hours of infection, two coexisting groups of cells were observed — one with clear detection of active viral expression (≥ 10 viral UMIs) and the other with transcript abundance at the level of noise (cf. control or mock infection). The observed bimodal distribution of viral transcripts implies the existence of an all-or-none switch that rapidly turns on viral expression (Fig. 1B). The proportion of cells with at least 10 viral UMIs increases from 0% to nearly 90% within 10 hours (Fig. 1C). Interestingly, the mRNA pool in each individual cell was mostly dominated by either host or viral transcriptome, with very few cells in intermediate states, forming a sharp decline in host-virus transcript ratio (Fig. 1D and fig. S2), indicating EhV massively transforms the cellular transcriptome by taking over almost the entire mRNA pool (fig. S3).

To further explore the variation among infected single cells, we applied the MetaCell analysis (*20*) based on viral gene expression, which divided the cells into seven groups (metacells) that defined the different phases of infection (Fig. 2A). Infected cells were spread along an infection continuum, reaching maximum heterogeneity at 6 hpi (Fig. 2B). In contrast, cells pre-treated with cycloheximide, an inhibitor of eukaryotic translation (*21*), were mainly localized in the initial phase (metacell 1) of infection at 6 hpi (Fig. 2B). This suggests that newly synthesized proteins are required for later viral transcription programs, but not for the initial phase. The continuum of cells allows us to map infection progression on a pseudotemporal scale between of 0 to 1 (Fig. 2A), providing a new dimension to determine infection states of individual cells within heterogeneous populations.

**Fig. 2.**
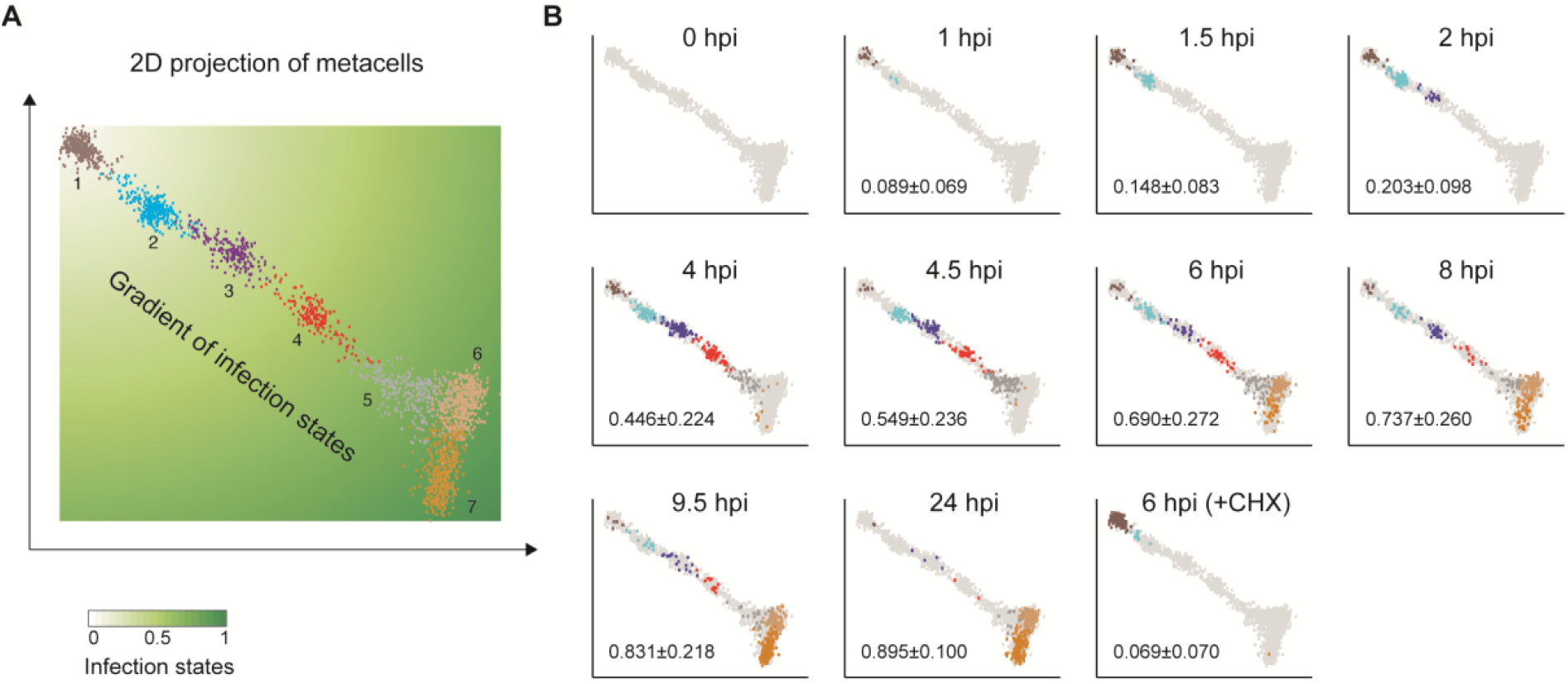
The continuum of infection states across individual cells. (**A**) 2D projection of 2,072 single infected cells (dots) with active viral expression (≥ 10 viral UMIs) that constitute seven metacells (in different colors). The cells form a continuum of infection states that can be quantified by the relative distance to the initial infection state. (**B**) Distribution of cells from each sampling time point along the infection continuum, with light gray dots representing cells from the other time points. Numbers are mean±SD of the infection states as defined in **A**. CHX: cycloheximide treatment before infection. The infection index (**A**) of an individual cell was calculated as the ratio of its distance from the upper-left origin to the distance between the origin and the lower-right end. A color gradient was painted based on the scale of the infection indices between 0 and 1.

By re-ordering cells based on the newly defined infection states, we found that viral genes were largely organized in five discrete kinetic classes of genes herein termed immediate-early (73 genes), early (136), early-late (49), late 1 (22) and late 2 (40), which formed partially overlapping sequential waves of expression (Fig. 3 and Table S2). The early class encompasses most of the genes involved in information processing and metabolism, including DNA replication (e.g., DNA polymerase delta subunit and proliferating cell nuclear antigen) and lipid metabolism such as the unique EhV-encoded sphingolipid biosynthesis (*13*, *22*–*24*) (Fig. 3C). The early genes also encode four RNA polymerase II subunits (RPB1, RPB2, RPB3, and RPB6), which probably enable transcription independent of the host machinery. In addition, two small RNA polymerase II subunits (RPB5 and RPB10) are encoded in the early-late class. The late 1 and late 2 gene products include the packaging ATPase for packing viral DNA into virions and the major capsid protein, the main component of the virion capsid, respectively. In addition, our proteomic analysis showed that most proteins encoded by the two late classes are integral components of EhV virions (Fig. 3C and Table S3).

**Fig. 3.**
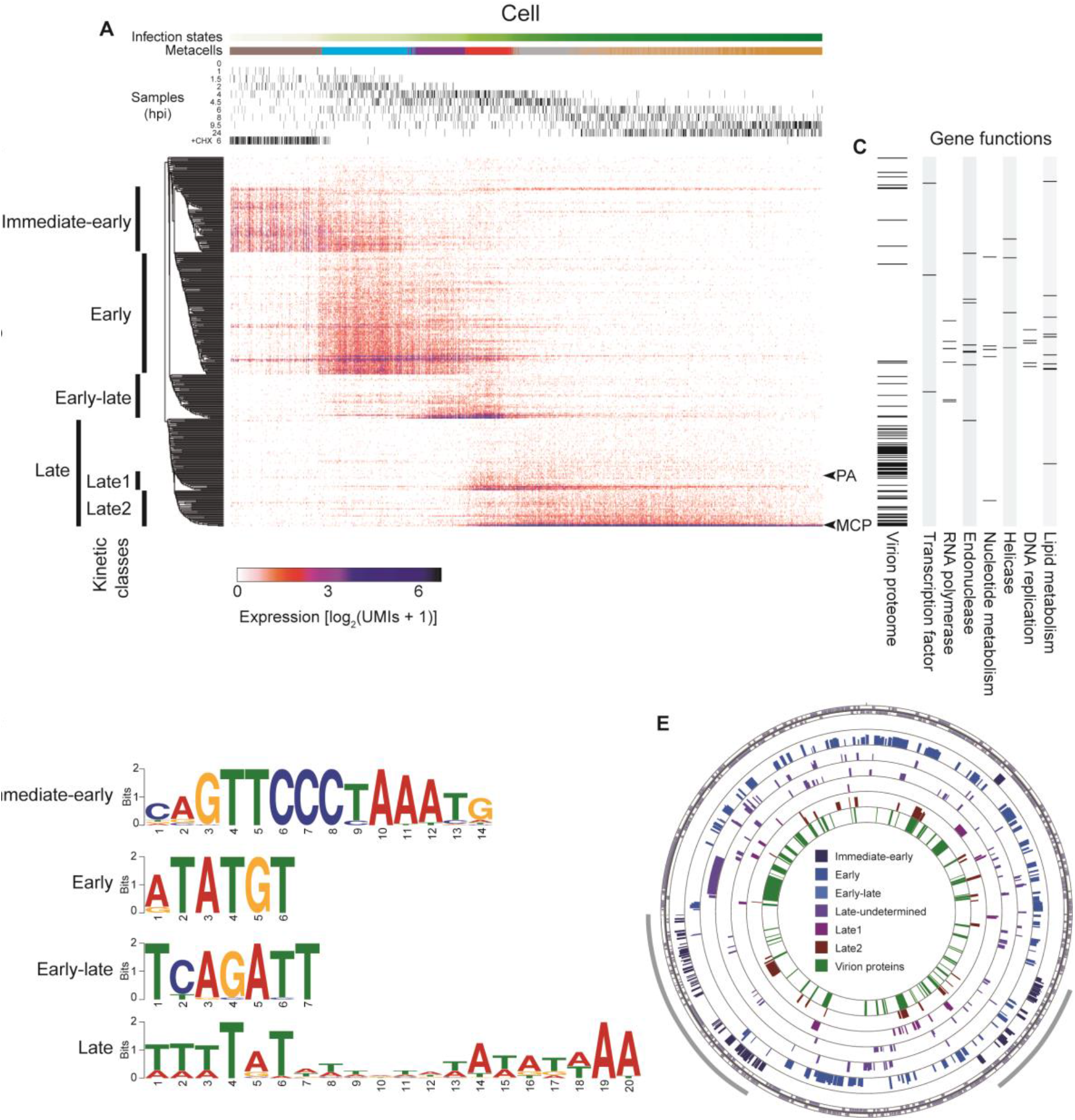
Viral kinetic classes, predicted promoter elements, and genomic organizations. (**A**) Cells (columns) are ordered by the pseudo-temporal scale (infection states) as defined in Fig. 2, with hours post infection (hpi) of each culture sample marked for each cell. (**B**) Viral genes (rows) are divided into kinetic classes based on the hierarchical clustering of their expression profiles across the individual cells. (**C**) Presence in the EhV201 virion proteome, as well as categories of predicted functions, are indicated for each gene. PA: packing ATPase; MCP: major capsid protein. (**D**) Enriched motifs in the promoter regions of viral genes in the newly defined kinetic classes. (**E**) Distribution of genes in each kinetic class (circle) on the assembled genome of EhV201. The outermost circle indicates the direction of transcription. The heights of genes in each kinetic class are proportional to their log_2_-transformed expression levels. The innermost circle marks the presence in the proteome of EhV201 virions. Immediate-early genes are mostly clustered in two genomic regions (gray arcs).

To characterize the regulatory mechanisms underlying the viral gene kinetic classes, we looked for enriched sequence motifs *de novo* in the promoter regions (±100 bp of the first base of the start codon) of genes belonging to different kinetic classes (Fig. 3D and Table S4). The most enriched motif in the promoter region of immediate-early genes ends mostly at the ATG start codon (positions 12-14) and was previously identified as a putative promoter element associated with initial expression in another EhV strain (*25*). We also revealed short, highly enriched elements, including a 6-bp motif around the ATG of early genes and a 7-bp one mostly upstream of the start codon in early-late genes. An AT-rich 20-bp motif was detected, with a degenerated center, usually immediately upstream of the start codon of late 1, late 2, and other late genes. We further found that these putative promoter elements are found to be conserved across EhV genomes (fig. S4), which share most of their gene contents (fig. S5 and Table S5). Intriguingly, the late motif bears striking similarity to the late promoter element of mimivirus, a distantly related NCLDV that infects amoebae, which is comprised by two AT-rich decamers separated by a degenerated tetramer and is also present in the genome of the mimivirus virophage (*11*). Although genes in most kinetic classes are scattered throughout the EhV201 genome, the immediate-early genes are mainly concentrated in two genomic regions (Fig. 3E), as also seen in the strain EhV86 (*25*). Such organization could facilitate the rapid activation of the immediate-early class once the host cell is infected, allowing transcription initiation of the entire cascade of the various kinetic classes. Taken together, our findings of newly defined viral kinetic classes are strongly supported by the discovery of potential regulatory mechanisms that are conserved across EhV genomes.

Finally, our single-cell dual RNA-seq approach allows quantification of not only transcripts encoded by the viral genome, but also respective host transcriptomes including those of the mitochondria and chloroplasts (Fig. 4). As depicted in Fig. 1, there was a sharp shift of the cellular transcriptome from host-encoded to virus-encoded transcripts. By mapping the expression levels across infection states, we found differential shutdown of nucleus-encoded and organelle-encoded genes (Fig. 4A). Whereas nuclear transcripts were rapidly shut down at the onset of active viral expression, levels of the mitochondrial and chloroplast transcripts were maintained. Moreover, the two types of organelles show distinct expression patterns, with mitochondrial transcripts persisting at lower, yet relatively stable levels throughout the infection progression and chloroplast transcript levels that were high in the beginning but then soon declined (Fig. 4B and 4C). The delayed shutdown of chloroplast-encoded genes suggests the need for active chloroplasts in early stages of infection, which is in agreement with the recent finding that photosynthetic electron transport remained active during the initial phase of viral infection (*26*). In contrast to chloroplasts, the mitochondrial transcripts were maintained throughout the infection progression, consistent with the high requirement for energy and redox equivalents at all stages of infection.

**Fig. 4.**
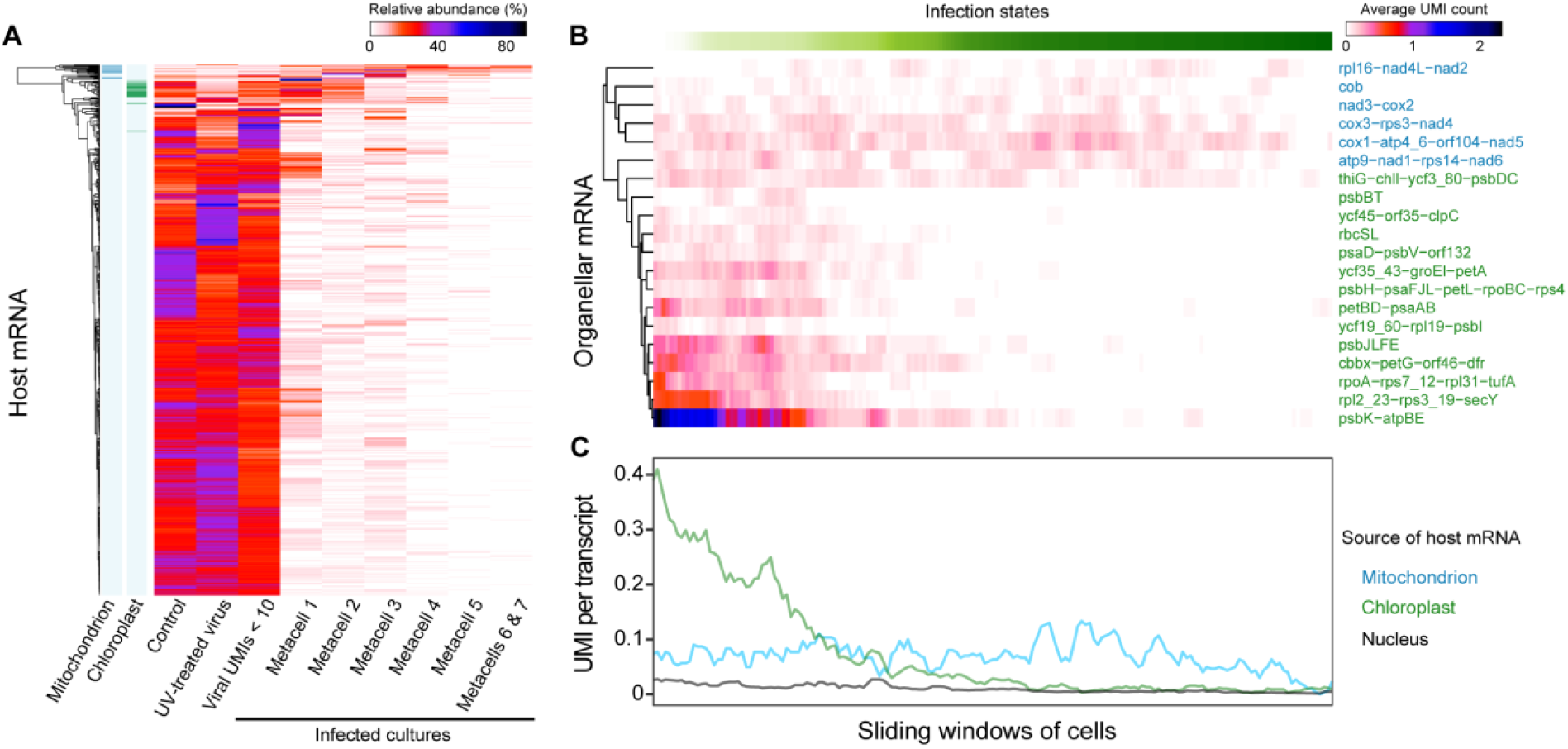
Highly resolved dynamics of virus-induced shutoff of host mRNA. (**A**) Levels of host transcripts across categories of cells. Single cells from cultures infected with EhV are divided based on overall viral UMI counts and metacell grouping (Fig. 2A). Protein-encoding genes with at least 100 UMIs were included. For each gene, the average UMI count was calculated for each category, and the sum of the average values of all categories was normalized to 100. Transcripts were then hierarchically clustered based on the normalized values. Organelle-encoded transcripts are indicated on the left. (**B** and **C**) Abundance of organellar transcripts along the infection progression. Cells with at least 10 viral UMIs are ordered according to the infection states as determined by the MetaCell analysis (Figs. 2 and 3). Sliding window averages were calculated for each transcript (B) or mean of all transcripts (C) with a window size of 50 cells and steps of 10.

Single-cell RNA-seq has proven to be a very powerful tool for comprehensive understanding of complex host-virus dynamics, disentangling cells from different sampling times and infection states. Using this approach, we dissected the transcriptomic dynamics and regulation of viral infection in a globally important microalga and showed rapid selective shutdown of host genes induced by an NCLDV. The comprehensive high-resolution mapping of infection states and kinetic classes allowed identification of conserved regulatory sequences which likely orchestrate the sequential expression programs during the viral life cycle. The applicability of scRNA-seq provides new resolution to track host-virus dynamics and holds great potential for *in situ* quantification of active viral infection in natural populations as well as identification of additional ecologically important host-virus systems. This in turn will facilitate the assessment of the impact of viruses on the function and composition of microbial food webs in the marine environment.

## Acknowledgements

We thank N. Stern-Ginossar, A. Nachshon, M. Shnayder, and E. Koh for help with MARS-seq and cycloheximide experiments; R. Avraham, D. Hoffman, S. H. Avivi, G. Rosenberg, A. Solomon, and Advanced Sequencing Technologies Unit of the Life Science Core Facilities for assistance with Illumina sequencing; Y. Levin and M. Kupervaser from the de Botton Institute for Protein Profiling of the Nancy and Stephen Grand Israel National Center for Personalized Medicine for assistance with proteomic analyses; and all Vardi lab members for discussion.

## Author contributions

C.K., U.S., S.R. and A.V. designed the study, with input from A.S.-P. and A.T.; U.S. and C.K. prepared the cell cultures; C.K. and U.S. performed MARS-seq with assistance of A.S.-P.; C.K. analyzed the MARS-seq data with assistance of A.S.-P. and A.T.; D.S. performed the virion proteome experiment; S.B.-D. analyzed the motifs and compared the EhV genomes, with input from C.K.; C.K. prepared the figures, with input from U.S., S.B.-D., S.R. and A.V.; C.K., S.R. and A.V. wrote the manuscript, with input from all authors; A.V. supervised the study.

## Data and materials availability

All data was deposited in Gene Expression Omnibus with the accession number GSE135429.

## Supplementary materials

### Materials and Methods

#### Algal culture and viral stock

The coccolithophore *Emiliania huxleyi* strain CCMP2090 was grown in K/2 (*27*) filtered seawater medium at 18 °C with an irradiance of 100 μmol photons m^−2^s^−1^ and a 16:8 h light-dark cycle. The viral strain EhV201 (*28*) was propagated on CCMP2090 culture until the culture lysed to clearance, and viral lysate was filtered using 0.45-μm PVDF Stericup-HV vacuum filtration system (Merck). We used the plaque assay as previously described (*29*) to quantify infectious virions, and filtered viral lysate generally contained ~10^8^ infectious virions per ml. Cultures at the exponential phase (~10^6^ cells ml^−1^) were used for viral infection at a high multiplicity of infection (5 infectious virions to 1 cell) at 2 h after the light cycle began. Algal cells and virion particles were enumerated using an Eclipse Flow Cytometer Analyzer (iCyt) during an infection time course (fig. S1). UV-inactivated virus was obtained by exposing viral lysate in a plastic petri dish to UV in a transilluminator system for 15 minutes. For inhibition of protein synthesis, cycloheximide was added to culture for a final concentration of 1 μg/mL immediately before viral infection.

#### Single-cell isolation by fluorescence-activated cell sorting

Single cells were isolated into individual wells in 384-well plates on a FACSAria II or III sorter (BD) using a 488-nm laser for excitation and a nozzle size of 100 μm. Cells were identified based on chlorophyll red autofluorescence (663–737 nm) and separated under the “Single Cell” mode for maximal purity.

#### Massively parallel single-cell RNA sequencing

The scRNA-seq method is based on the MARS-seq2.0 protocol (*19*). Before sorting, each 384-well plate was prepared using a Bravo liquid handling platform (Agilent) to transfer into each well 2 μl lysis buffer containing 2 or 4 nM reverse transcription (RT) primers with cellular and molecular (UMI) barcodes. Plates with sorted cells were immediately frozen on dry ice and later stored at −80 °C. After thawing, the plates were heated to 95 °C for 3 min to evaporate the lysis buffer, cooled, added RT reagent mix, and placed in a Labcycler (SensoQuest) for RT reaction. The unused primers were then degraded with Exonuclease I (NEB). Complementary DNA from each set of wells with 384 (one pool per plate) or 192 (two pools per plate) unique cellular barcodes were then combined into pools, which underwent second strand synthesis and *in vitro* transcription to amplify the sequences linearly. The RNA products were fragmented and ligated with single-stranded DNA adapters containing pool-specific barcodes. A second round of RT was carried out, followed by polymerase chain reaction with primers containing Illumina adapters to construct DNA libraries for paired-end sequencing on a NextSeq or MiniSeq machine (Illumina).

#### Read processing and mapping to reference sequences

The fastq files were processed using the analytical pipeline of MARS-seq2.0 (*19*) which mapped the reads to viral and host reference sequences and demultiplexed them based on the pool, cellular, and molecular barcodes. At least 4 reads per UMI were obtained per cell. For the virus, the predicted coding sequences (CDS) in the EhV201 genome sequence (JF974311) (*30*) were used as reference. For the host, an integrated transcriptome reference of *E. huxleyi* (*31*) was used, where both forward and reverse (labeled with “_rev”) strands of each sequence were included for mapping because the CDS directionality was often unknown. In addition to nuclear sequences, the transcriptome reference contains chloroplast and mitochondrial transcripts, which were identified by BLAST (*32*) searches against the respective organellar genome sequences (*33*, *34*).

#### Viral and host transcript abundance

To avoid empty wells or those with low transcript capture during library preparation, wells with fewer than 10 UMIs were removed for all downstream analyses. Wells with more than 1,000 viral transcripts were also removed to prevent wells with doublet or multiplet cells from confounding single-cell infection state analyses. The vast majority (> 99%) of control or infected *E. huxleyi* cells had fewer than 1,000 total UMIs, which reflects their low RNA content due to the small cell size (3-5 μm in diameter for naked cells (*35*)).

#### Cliff diagrams of viral and host mRNA

A cliff diagram was plotted for each time point post infection (Fig. 1D). The total viral mRNA transcript counts were calculated by summing up all UMIs assigned to virus-encoded CDS. The total host mRNA counts were all host UMIs except for those assigned to the three transcript sequences corresponding to nuclear, mitochondrial, and chloroplast ribosomal RNA genes. For each sampling time point, the cells were ordered by the ratio of host to all (host+virus) mRNA from high to low. Cells with fewer than 20 total UMIs were excluded from the plots to avoid sampling biases.

#### Doublet assay and in silico simulations

The EhV201 viral genome has a much lower GC content (40.46%) (*30*) than the *E. huxleyi* host genome (65%) (*36*). This difference in GC content and other factors might bias the capture and amplification of viral and host transcripts during MARS-seq and might lead to the either-or dominance of host and viral mRNA (Fig. 1D). To test if the presence of viral mRNA has effects on the quantification of host mRNA transcripts and vice versa, we conducted a doublet assay. Half of the doublet assay plate consisted of single cells sorted from infected culture at 8-hpi and the other half pate consisted of two cells per well of which one cell was sorted from infected culture at 8 hpi and the other cells from control culture (Table S1). Cliff diagrams were then plotted for each half of the plate (fig. S2A-B). If the viral mRNA does not interfere with host mRNA quantification, wells with cells isolated from both the infected and the control cultures should show host and viral mRNA UMI counts that are additive, as if the two cellular transcriptomes were sequenced separately and merged together. To simulate this *in silico*, we randomly sampled sequenced single-cell transcriptomes from the control culture plate with the closest timepoint to 8 hpi (Table S1) using the *sample* function in R, and they were combined one-by-one with those from the 8-hpi half plate with single cells. Cliff diagrams were plotted for each of the eight simulations (fig. S2C-J).

#### MetaCell analysis of infected cells

The MetaCell package is an unbiased approach for cell grouping based on single-cell expression profiles without enforcing any global structure (*20*). To account for the small number of UMIs per cell and to use more viral genes for grouping infected cells, we employed more inclusive criteria for gene marker selection (T_tot=20, T_top3=1, T_szcor=−0.01, T_vm=0.2, and T_niche=0.05). A total of 179 marker genes were used constructing *k*-nearest neighbours graphs with K=150, followed by co-clustering with bootstrapping based on 1,000 iterations of resampling 75% of the cells and an approximated target minimum metacell size of 80. Unbalanced edges were filtered with K=40 and alpha=3.

After plotting single cells in a 2D projection and identifying the directionality of infection progression (Fig. 2), we quantified the infection states of individual cells based the relative distance to the beginning of the progression (i.e., upper-left). The *ad hoc* origin was defined as having x- and y-coordinates of the leftmost and the highest cells, respectively, and the *ad hoc* end as having those of the rightmost and the lowest cells, respectively. The infection index of an individual cell was calculated as the ratio of its distance from the origin to the distance between the origin and the end. A color gradient was painted based on the scale of the infection indices between 0 and 1.

#### Hierarchical clustering of genes

We used hierarchical clustering to group viral genes with similar expression patterns across infected cells. Viral genes with fewer than 20 UMIs in total across all the cells with active viral expression were excluded to avoid spurious clusters or groupings that are not well supported. The pheatmap package in R was used for hierarchical clustering (hclust) of UMI counts in log scale, using Pearson correlation as distance measure and the “average” agglomeration method. The cluster tree of viral genes was manually curated by branch swapping using Archaeopteryx (*37*) to order the genes by kinetic classes and by expression levels.

#### Annotations of EhV201 genome

A comprehensive annotation table was prepared for EhV201 genes by ordering them according to the manually curated cluster tree (Fig. 3B). The sequence descriptions were based on the GenBank record (JF974311), with modifications based on relevant publications (*38*, *39*), BLAST searches against the nr database (*40*), and the Nucleo-Cytoplasmic Virus Orthologous Groups (NCVOGs) (*2*).

#### Virion proteome composition

Purified EhV201 virion samples were lysed using an 8 M urea buffer and subjected to tryptic digestion followed by LC-MS analysis on a Q Exactive HF instrument (Thermo). We used Byonic version 2.10.5 (Protein Metrics) as the software to process the data and to search against the viral CDS database (Table S3). In total 76 viral CDSs had at least 10 peptide-spectrum matches and were shown as present in virion proteome in Figure 3.

#### Sequence motif analyses of viral genomes

A 200-bp region surrounding the predicted start codon ATG (A of ATG is base 101) was extracted for each CDS in the EhV201 genome using the Extract Genomic DNA tool of Galaxy (*41*). Promoter motif discovery was performed on the positive strand of these regions for genes in different kinetic classes using MEME version 5.0.5 (*42*), Genomatix Genome Analyzer CoreSearch (Intrexon Bioinformatics), and Improbizer (https://users.soe.ucsc.edu/~kent/improbizer/improbizer.html), with the number of motifs to detect set to 4. While all three programs agreed on the motifs detected, the results of the MEME analyses are used in this study, where the most common motifs of lengths 6-20 are shown (Fig. 3D, fig. S4, and Table S4). To test if the short 6-bp sequence motif in early genes is significantly enriched, a differential analysis was performed for early genes against all late (late 1, late 2 and late-undetermined combined) using DREME (*43*) (Table S4).

#### Circular visualization of viral genes

The UMI counts of individual viral genes were summed over all cells showing active viral expression (≥ 10 viral UMIs). The log_2_ values of these total UMI counts were visualized using CCT (*44*), with a ring for each kinetic class and the height proportional to the log-transformed expression value. Another ring shows the presence/absence of each gene product in the virion proteome (Fig. 3E).

#### Comparative analyses of other EhV genomes

The best BLAST hits in the CCT analyses were used to find the homologous proteins of the various kinetic classes in other EhV genomes. DNA surrounding the ATG of those genes was extracted for each strain with the GetFastaBed tool in Galaxy (*41*). Promoter analyses were run on the sequences of the promoter regions of three additional strains (EhV207, EhV86 and EhV202) individually as described above and with all four together (fig. S4 and Table S4). These three strains were chosen as representatives of three major clades of EhV strains (*45*). A CCT plot was drawn to show the sequence similarity between EhV201 and all 12 other EhV strains with whole genome sequences (fig. S5).

#### Nuclear and organellar transcripts

Host gene expression patterns were analyzed for different categories of single cells, including control culture, mock infected culture, infected culture with fewer than 10 viral UMIs, and different metacells of infected cells with at least 10 viral UMIs. Transcript sequences with fewer than 100 total UMIs across all the cells were excluded to avoid lowly expressed ones, leaving 720 unique transcripts in total (Fig. 4A). The average UMI count was calculated for each transcript in each category, and the sum of the average values of all categories was normalized to 100. Hierarchical clustering of host transcripts was done using pheatmap as described above based on the normalized values. The sliding windows (window size = 50 cells, step = 10 cells) for organellar transcripts were generated using the rollapply function of the zoo package in R.

**Fig. S1.**
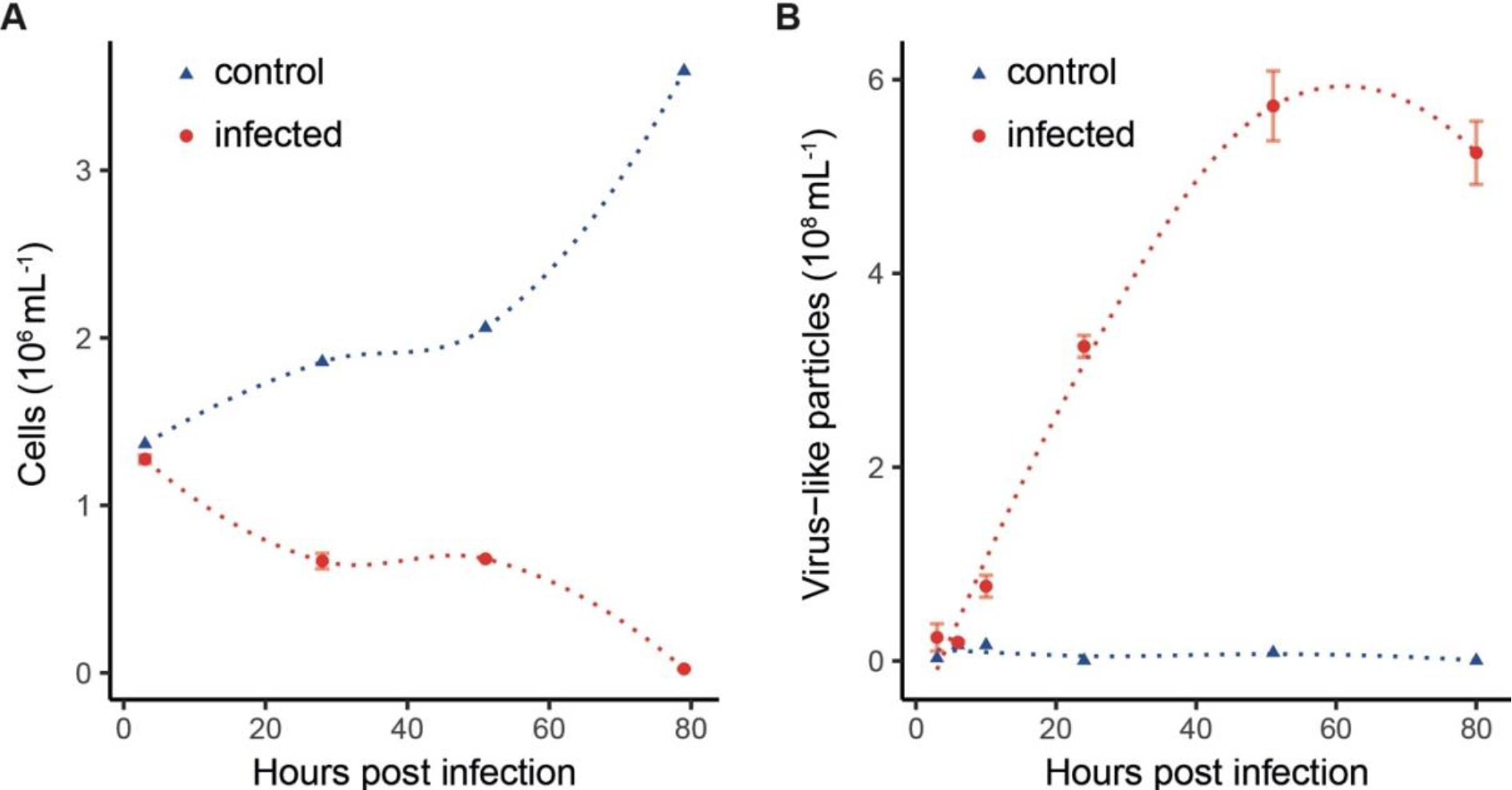
Abundance of *E. huxleyi* cells and extracellular EhV201 virions during a time course of infection. (**A**) Numbers of *E. huxleyi* cells in control and infected cultures. (**B**) Numbers of virus-like particles cells in control and infected cultures. A line was plotted for each treatment using the locally estimated scatterplot smoothing method (span = 1) implemented in the ggplot geom_smooth function in R. The infected culture had two biological replicates, and mean ± SD is shown.

**Fig. S2.**
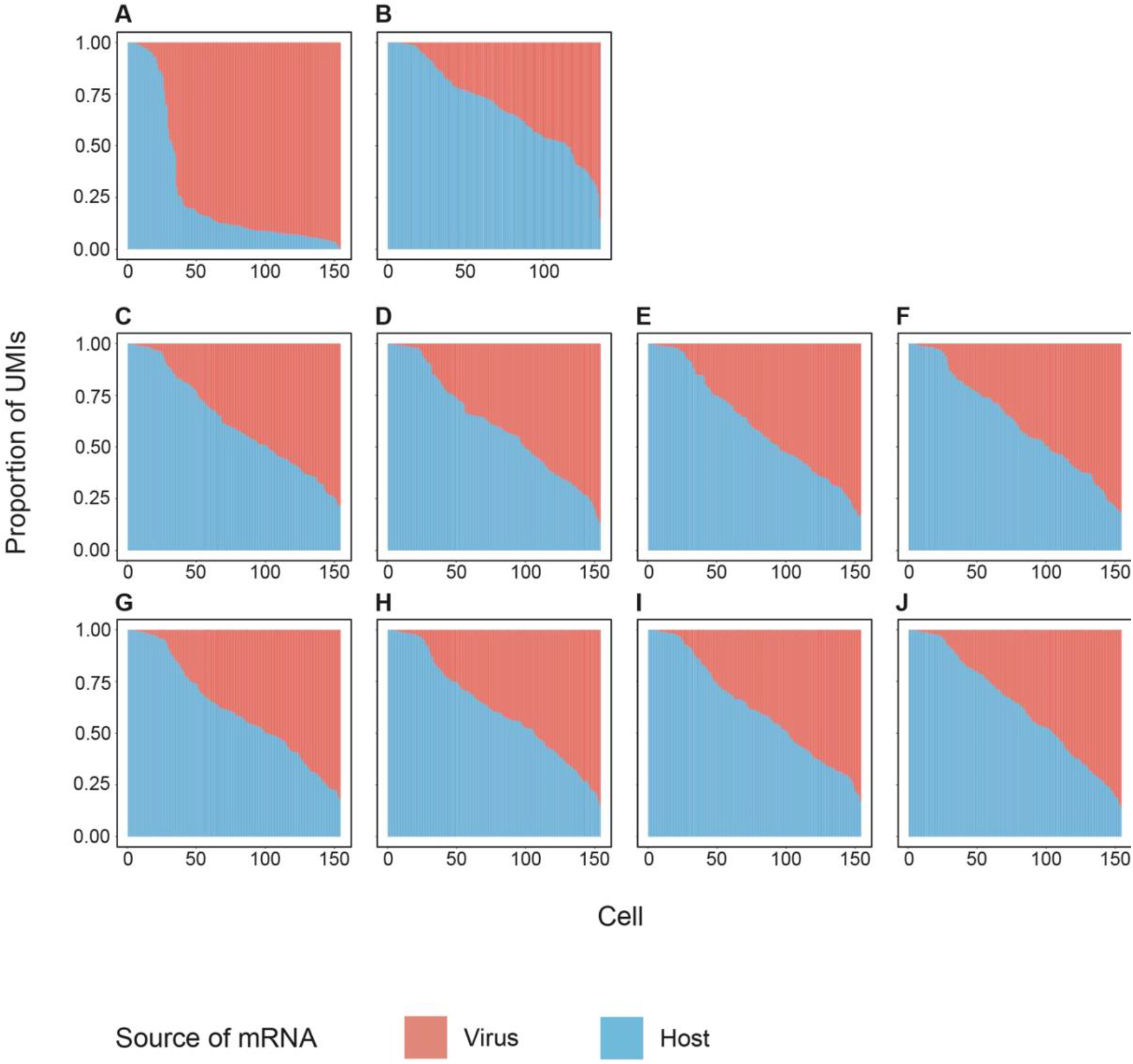
The presence of viral mRNA does not affect the capture and sequencing of host mRNA transcripts by MARS-seq. To test if the shutdown of host mRNA (Fig. 1D) is not due to a possible bias of MARS-seq toward viral transcripts given the differences in GC% content and other properties between viral and host genes, a doublet assay was conducted. Cells at 8 hpi were sorted and viral and host transcript quantifications were compared between wells with a single cell from infected culture (half a 384-well plate) and wells with one from infected and one from control cultures (the other half of the plate) (Table S1). Cliff diagrams are used to represent viral and host proportions of the mRNA pool in single cells (Fig. 1D). (**A**) Wells with one cell sorted from the infected culture at 8 hpi. (**B**) Wells with one cell sorted from the infected culture and one from the control culture at 8 hpi. (**C**–**J**) Cliff diagrams based on four *in silico* simulations, where the same number of cells from control plate 10 hpi were sampled and randomly paired with the single cells from the infected culture at 8 hpi (i.e., **A**), show patterns similar to the experimental results (**B**). This observation indicates that the presence of viral molecules did not affect the quantification of host transcripts and that the absolute counts of these transcripts are additive. For the doublet experiment (**B**) or simulations (**C**–**J**), a “cliff” (sharp change in mRNA ratios) is absent; instead, a “slope” (gradual change) is observed because of random pairings of infected and control culture cells of different cell size and transcript content.

**Fig. S3.**
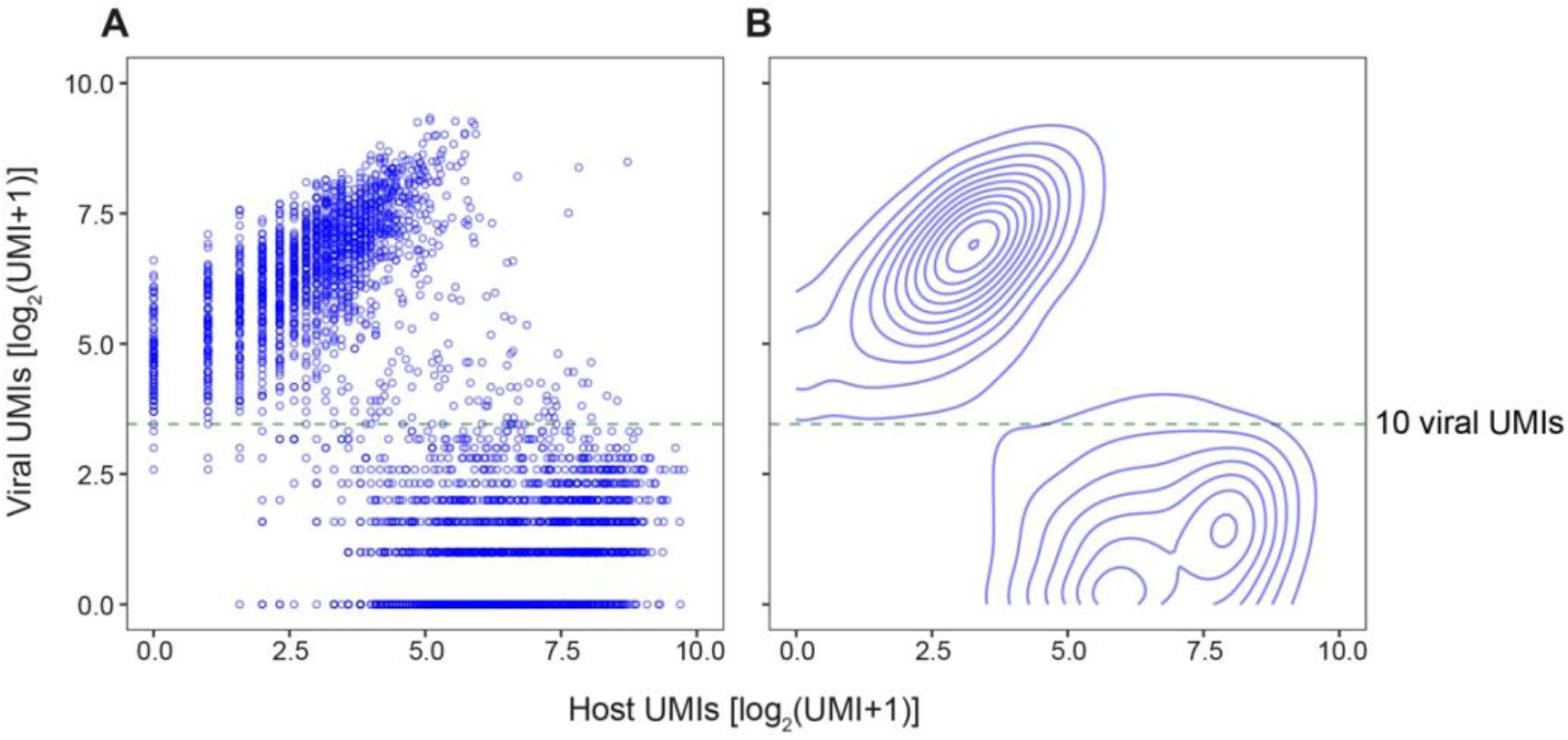
Abundance of viral and host mRNA across single cells in EhV-infected culture of *E. huxleyi*. (**A**) For each cell from infected samples and with total UMIs between 20 and 1,000, the numbers of viral and host mRNA UMIs are plotted. (**B**) A contour plot of **A** based on 2D kernel density estimation using the geom_density_2d function of ggplot2. A threshold of 10 viral UMIs filters out wells with background or noise levels of viral UMIs and separates cells with and without host shutdown.

**Fig. S4.**
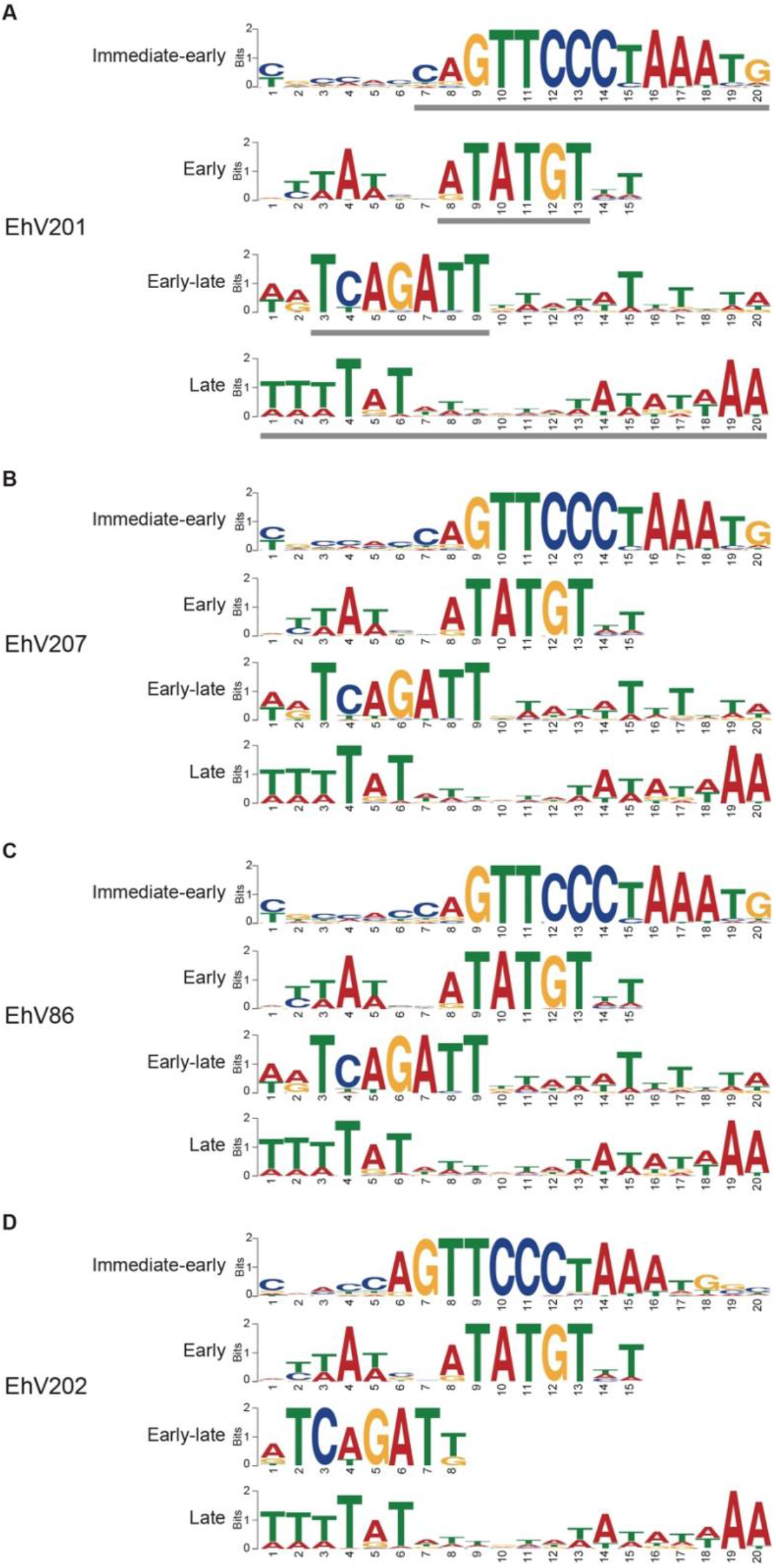
Promoter elements specific to EhV201 kinetic classes are conserved across EhV strains. (**A**) Enriched sequence motifs (6-20 bp) found in the promoter regions (±100 bp of the first base of the start codon) of genes in different kinetic classes of EhV201. The gray bars indicate the highly enriched motifs shown in Fig. 3D. (**B**-**D**) Enriched sequence motifs in the promoter regions of genes in three other EhV strains that are homologous to the EhV201 genes in different kinetic classes. The three strains represent the three major clades of EhV strains (*45*) (Table S4) and are ordered with increasing divergence from EhV201. The numbers on the x-axis indicate the positions within each motif.

**Fig. S5.**
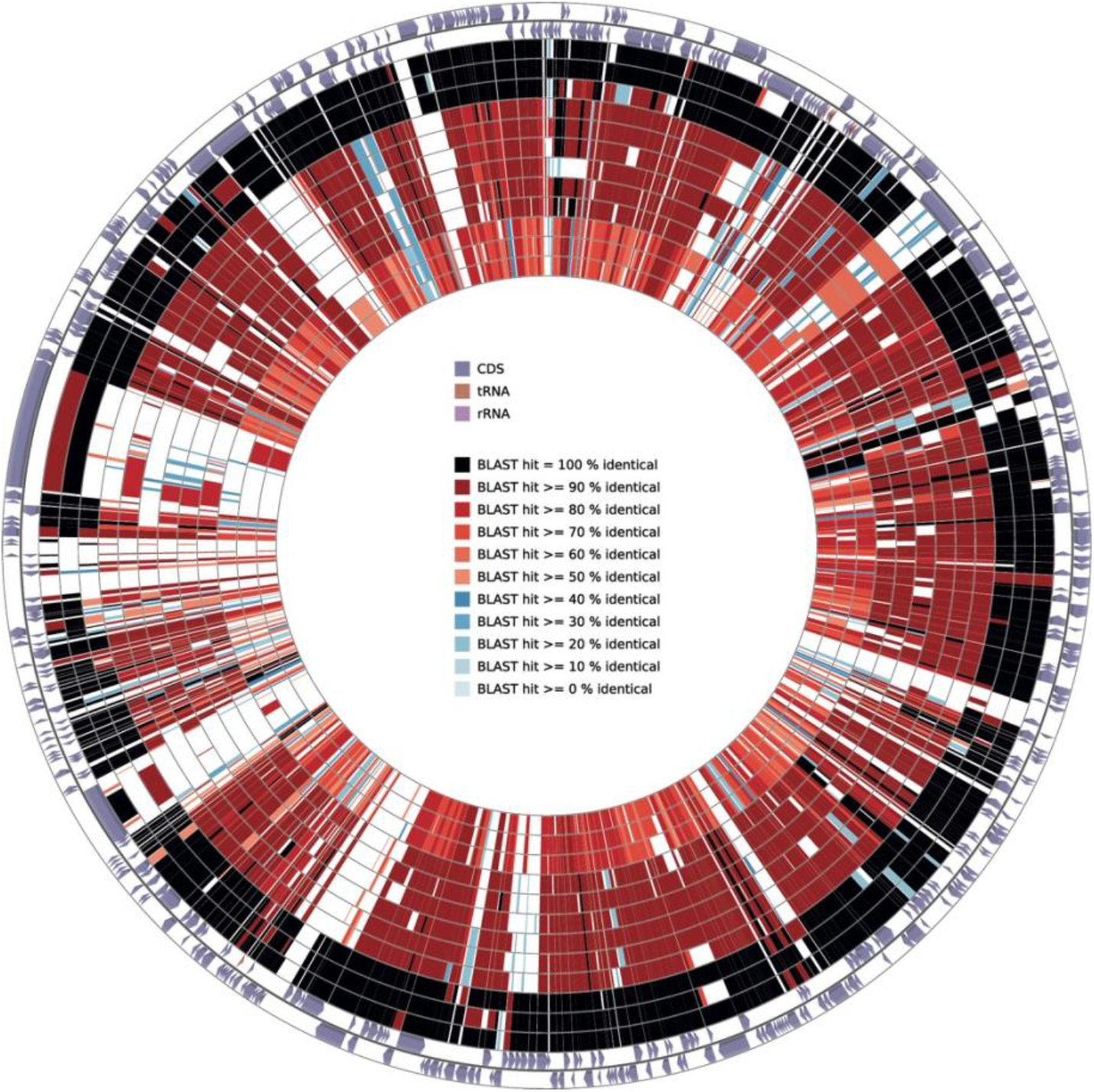
The EhV201 genes are largely conserved in 12 other EhV genomes. In this similarity circular plot, the two outermost circles indicates the direction of transcription and types of genes. The other circles show the BLAST similarity of protein sequences between EhV201 and the following strains (from outer to inner): EhV207, EhV203, EhV208, EhV88, EhV86, EhV84, EhV99B1, EhV164, EhV145, EhV202, EhV156, and EhV18. See also Table S5.

